# Cortical sensorimotor activity in the execution and suppression of discrete and rhythmic movements

**DOI:** 10.1101/2021.09.30.462656

**Authors:** Mario Hervault, Pier–Giorgio Zanone, Jean–Christophe Buisson, Raoul Huys

## Abstract

Although the engagement of sensorimotor cortices in movement is well documented, the functional relevance of brain activity patterns remains ambiguous. Especially, the cortical engagement specific to the pre-, within-, and post-movement periods is poorly understood. The present study addressed this issue by examining sensorimotor EEG activity during the performance as well as STOP-signal cued suppression of movements pertaining to two distinct classes, namely, discrete vs. ongoing rhythmic movements. Our findings indicate that the lateralized readiness potential (LRP), which is classically used as a marker of pre-movement processing, indexes multiple pre- and in-movement-related brain dynamics in a movement-class dependent fashion. In- and post-movement event-related (de)synchronization (ERD/ERS) observed in the Mu (8-13 Hz) and Beta (15-30 Hz) frequency ranges were associated with estimated brain sources in both motor and somatosensory cortical areas. Notwithstanding, Beta ERS occurred earlier following cancelled than actually performed movements. In contrast, Mu power did not vary. Whereas Beta power may reflect the evaluation of the sensory predicted outcome, Mu power might engage in linking perception to action. Additionally, the rhythmic movement forced stop (only) showed a post-movement Mu/Beta rebound, which might reflect an active “clearing-out” of the motor plan and its feedback-based online control. Overall, the present study supports the notion that sensorimotor EEG modulations are key markers to investigate control or executive processes, here initiation and inhibition, which are exerted when performing distinct movement classes.

## Introduction

It has long been known that when performing a voluntary action, cortical sensorimotor areas are engaged in movement planning, execution and online control ^1^. Most corresponding accumulated knowledge has been acquired in the context of the generation of discrete movement, which constitute an important, but not sole class of movements that humans can perform ^2^. Consequently, two aspects of action control and its neural sensorimotor underpinnings are strongly under-represented. On the one hand, we know little about cortical sensorimotor engagement related to movement suppression, even though both movement generation *and* suppression are commonplace in our interaction with the environment ^3^. On the other hand, previous investigations of neural activity when suppressing movements have focused exclusively on short-lived discrete movements and have then ignored the case of *ongoing-rhythmic* movement suppression, which is also crucial in action control ^4–7^. The few studies at hand on sensorimotor activity related to action suppression have dealt with prepared discrete movements ^8–11^, discrete movements sequence ^12,13^ or isometric force exertion ^14,15^. Kinematically, discrete actions are delimited by moments without movement (i.e., with zero velocity and acceleration), such as grasping an object. In contrast, continuous actions, such as walking, lack recognizable endpoints and are typically considered rhythmic if they constitute (periodic) repetitions of particular events ^2^. Motor control encompasses both action classes, which differ not only regarding their kinematics ^16^ but also in terms of movement dynamics and control processes ^17,18^, as well as of corresponding brain engagement ^19^. Indeed, the neural structures associated with controlling discrete and rhythmic actions differ considerably ^19–21^, due to different timing and initiation mechanisms ^17,20^. Additionally, integrating in- and post-movement sensory information shows distinct dynamics between discrete and rhythmic action classes ^22,23^, which may involve open- and closed-loop control, respectively. As sensorimotor EEG activity has been linked to movement-related sensory integration in the framework of forward internal models of motor control (see below), its investigation and comparison in both movement classes appears to be crucial.

The present study aims to help providing a more complete picture of the cortical sensorimotor activity underlying action control through the study of both the performance *and* suppression of movements belonging to two fundamentally distinct classes, discrete *and* rhythmic movements. EEG activity over sensorimotor areas was analyzed in terms of the lateralized readiness potential (LRP) and event-related (de)synchronization (ERS/D) of Mu (8 - 13 Hz) and Beta (15 - 30 Hz) cortical oscillations. In addition, a second objective was to provide new insights into understanding the functional relevance of these movement-related neural sensorimotor activities with regard to action executive control.

Prior work has established standard non-invasive methods to explore movement-related brain activity. When recording scalp EEG, the LRP is believed to reflect the central response preparation within the primary motor cortex (M1) that control the movement ^24^. As for brain oscillations, a well-defined pattern of activity has been described during and after movement execution in Mu and Beta rhythms. This pattern is characterized by an ERD associated with the movement’s execution, followed by an ERS subsequent to the movement stop ^25^. This ERD/ERS pattern has been recorded over sensorimotor areas for several (contrasting) movement conditions, including self-paced and stimulus-triggered movements ^26,27^, real and imagined movements ^28^, as well as discrete short responses and lasting rhythmic movements ^29,30^. Especially, the cortical ERD/ERS dynamics were clearly observed for each movement cycle in the case of low-frequency movement repetition (< 1 Hz), that is, when the repetition was most likely due to a concatenation of discrete movements. In contrast, it transformed into a sustained ERD during higher-frequency movement repetition, that is, when the movements were truly rhythmic ^30–32^.

Despite the large number of studies reporting these movement-related neurophysiological modulations, their functional relevance remains debated. The LRP is thought to reflect the pre-movement M1 engagement as a final pathway for the central generation of movement, that is, the downstream specification of commands to the peripheral motor structures ^33^. Accordingly, LRP is massively used as an index of movement initiation when triggering discrete movement across multiple simple and choice reaction time tasks ^34,35^. In this context, LRP may follow a fixed-threshold dynamics, that is, the reaching a threshold activation amplitude determines whether the response will be triggered or not ^36,37^. Based on the assumption that the reach of this threshold discriminates successfully from failed cancellations of a prepared discrete movement ^33^, LRP has become a popular tool for investigating discrete action inhibition ^38–41^. When performing a continuous action, an external signal may indicate the performer to speed up ^42^, continue ^43^ or stop ^6,12,42^ the ongoing action. In such cases, a new command specification might engage in the building up of the motor activity. However, the purported assignment of LRP to pre-movement processing has led to its dereliction for investigating the voluntary modulation or suppression of an ongoing rhythmic movement. Indeed, the very possibility of an LRP reduction has been ignored by the few studies exploring rhythmic movement stopping ^7,43^.

The Mu/Beta ERD reflects the desynchronization of an ensemble of cortical neurons over sensorimotor brain areas. In contrast, the post-movement Mu/Beta ERS reflects its neural resynchronization ^44^. The Mu/Beta activity has been initially suggested to echo a cortical idling state during “mental inactivity” ^45^ or a “status quo” in maintaining the current sensorimotor or cognitive state ^46^. Although Mu and Beta tend to follow a similar pattern of activity and can be mapped to a single dipole due to an overlap in their cortical sources, recent evidence showed that they index distinct neurological functions ^47^. These functions, which are still debated, have been proposed in the framework of forward internal models of motor control ^48^, in which the sensory consequences of movement are predicted (through forward models) and compared to the actual sensory outcome. Indeed, the Mu rhythm has been considered as an alpha-like oscillation engaged in a “diffuse and distributed alpha system”, in reference to the multiple ∼10 Hz rhythms originating from independent brain sources ^49^. Within this broad alpha system, the Mu rhythm might reflect a perception-to-action translation ^47,50^. Accordingly, Mu synchronicity occurs when visual and auditory representations are converted into action-based representations. The potential distinction between sub-frequencies bands ^47^ and the Mu involvement in inverse models ^51^ is still examined. At any rate, the Mu rhythm is generally viewed as a correlate of the reciprocal interaction between motor and sensory cortices, this interaction being crucial in the internal models controlling the action.

According to recent reviews ^47,52^, the Beta ERD reflects movement preparation, including the adjustments of motor commands and the anticipation of errors ^53^. The Beta ERD modulation by movement uncertainty ^54^ also suggests that it plays a role in predicting the sensory consequences of the action. The observation of an above-baseline ERS following movement, known as the post-movement Beta rebound (PMBR), led to multiple hypotheses. Beta oscillations could reflect the post-movement processing of sensory reafference ^55^. Indeed, the occurrence of PMBR after passive movements ^56^ or when accompanying peripheral nerve stimulation ^57^ is consistent with the idea that PMBR originates in sensory feedback to the motor cortices. More specifically, the PMBR was proposed to index the integration of sensory feedback to evaluate movement outcome, with any deviation from the forward-predicted outcome leading to an update of the motor plan ^47^. Alternatively, PMBR could reflect the active inhibition of the motor cortex to terminate a movement ^58^. The observation of a single PMBR following a sequence of discrete movements ^13,59^ and its association to movement parameters such as accuracy, variability, and rate of force development ^60,61^ have been taken as an argument for its involvement in the active inhibition of the motor cortex following movement termination.

All in all, multiple interpretations have been put forth to explain neural sensorimotor activity before, during and after a movement. Additionally, in relation to the ERD/ERS pattern, the brain activation found over both pre- (motor) and post-Rolandic (somatosensory) areas ^50,52,62^ contributes to blur the numerous functional hypotheses. Still, experiments requiring both initiation and suppression of movement have tried to provide new insight into the functionality of the sensorimotor ERD/ERS by showing that its occurrence depends on whether a movement is actually performed versus withheld ^11^. The cortical activity also differed between normal movement completion and forced suppression ^12^ and between quick and slow movement termination ^14^. However, the characterization of the movement-related sensorimotor activity suffers from large variation in the task parameters employed across studies (e.g., task duration and movement amplitude), which alters the corresponding neural activity, and has hampered the establishment of convincing functional interpretations ^15^.

To complement our understanding of the movement-related neural sensorimotor activity, the present study examined EEG activity when performing a movement and suppressing it. EEG was recorded in the context of two fundamental classes of movement: discrete and rhythmic ones. Using a graphic tablet, we asked participants to initiate a discrete movement after a GO stimulus and pursue a rhythmic movement after a CONTINUE stimulus. Infrequently, a STOP signal following the primary stimulus indicated participants to cancel the prepared-discrete movement or to stop the ongoing-rhythmic one. Firstly, in line with the interpretation of LRP as a sign of movement preparation, we hypothesized its large amplitude following a GO stimulus to contrast with its absence following a CONTINUE stimulus, and the STOP signal occurrence to reduce its amplitude in the discrete experiment only. Secondly, following the assumption that Mu and Beta rhythms encode reciprocal interactions between motor and sensory cortices to enable monitoring of movement, we expected to observe a sustained Mu ERD during ongoing rhythmic movement ^30^, reflecting the closed-loop processing of sensory information in the CONTINUE condition, and it to be aborted by movement suppression in the STOP condition. In contrast, we expected to indifferently observe a transient Mu ERD/ERS in discrete completed, successfully cancelled, and unsuccessfully cancelled actions, as the movement is controlled in an open-loop fashion, and to observe a transient and sustained Beta ERD, reflecting motor activation, in the discrete and rhythmic condition, respectively. Third, we anticipated a PMBR, reflecting the post-movement sensory “check”, to be visible after movement suppression in the rhythmic STOP ^14^ and the discrete conditions, with differences between the discrete completed, successfully cancelled, and unsuccessfully cancelled actions, for the movement outcome differs in each case ^11^.

## Method

### Participants

Fifteen healthy individuals (9 males, mean age 25 years, SD = 2.2) served as voluntary participants. All were right-handed, as assessed by the Edinburgh Handedness Inventory ^63^, and had a normal or corrected-to-normal vision. None of the participants reported a history of psychiatric or neurological disorders. The study was conducted with the informed consent of all participants according to the principles stated in the Declaration of Helsinki, and the procedures were approved by the local research ethics committee (Comité de Protection des Personnes Sud-Ouest et Outre-Mer II; ID-RCB: 2020-A03215-34).

### Procedures

#### Experimental procedures

Participants performed two experiments that have been previously described ^43^, and for which details are provided in Appendix A. Briefly, both experiments required participants to perform voluntary right-hand movements on a graphic tablet using a stylus. In both experiments, the participants completed one practice block and 30 experimental blocks, each consisting of 20 trials. In the first experiment, visual GO stimuli called for the quick initiation of discrete-swipe movements (GO_D_ condition). Following the primary GO stimulus, a STOP signal was presented infrequently (in 25 % of trials, STOP_D_ condition), indicating the participants to cancel the prepared movement, leading to successful-STOP_D_ or fail-STOP_D_ trials. The experiment was designed following the recent guideline for stop-signal tasks ^64^. In the second experiment, participants executed self-paced rhythmic movements; a visual CONTINUE stimulus called for the continuation of a rhythmic movement (CONTINUE condition). As in the first (discrete) experiment, infrequently (in 25 % of trials, STOP_R_ condition), a STOP signal followed the primary CONTINUE stimulus to order participants to stop the ongoing movement quickly. Following such STOP trials, a rhythmic GO_R_ trial was added to reengage participants in the rhythmic movement. In these GO_R_ trials, participants were instructed to transit from a static position to an oscillating movement as soon as the GO stimulus (green or blue) was presented. In both the discrete and rhythmic experiment, the minimal delay between two trials was 3500ms and the primary stimulus occurrence varied randomly in a 500 ms window. As such, the two experiments are close in design in terms of the stimuli properties and the effectors engaged in the movement production; their main difference consisted in the movement type to perform and stop, namely prepared-discrete versus ongoing-rhythmic movements.

#### EEG recording and preprocessing

Scalp EEG was recorded using an ActiveTwo system (BioSemi Instrumentation, 64 electrodes) with a sampling rate of 2048 Hz. The EEG electrodes were cautiously positioned based on four anatomical landmarks (i.e., nasion, inion, and preauricular points) in accordance with the 5 % 10/20 international system ^65^. Additional electrodes were placed below and above each eye. The data were online referenced to the BioSemi CMS-DRL reference. All offsets from the reference were kept below 15 mV. The EEG data were filtered online with a frequency bandpass filter of 0.5-150 Hz. The participant’s arm was fixed on the table to restrain the movement to wrist articulation and avoid muscular noise in the EEG signal due to substantial contraction of the biceps and deltoid muscles. Continuous EEG data were imported and preprocessed in bespoke scripts using functions from the EEGLAB Matlab plugin ^66^ :

- Visual inspection was used to remove channels with prominent artifacts in the continuous EEG.
- The EEG data were then re-referenced to a common average.
- The data were partitioned into epochs of 3 s (locked to the primary stimulus onset; −1000 ms to 2000 ms).
- Those epochs containing values exceeding the average across the data segments by 5 SD were rejected.
- Scalp EEG data typically represent a mixture of activities originating from brain sources that are not separable based on channel data solely. Independent component analysis (ICA) ^67^ can be applied to identify statistically independent signal components (ICs) spatially filtered from the 64 channels data. An ICA was applied to continuous EEG data (concatenation of the EEG epochs) to identify 63 neural ICs contributing to the observed scalp data. Using the ICLABEL classifier ^68^ over the 30 first ICs, components with less than 10% chance to account for neural activity were considered as artifacts, and removed from the EEG data structure, thus removing their contributions to the observed EEG. The rejection was systematically verified by visual inspection of component properties (time series, spectra, topography) according to ICLABEL guidelines ^68^.

Across all participants, these procedures led to the omission of 8.6 % of the STOP trials in the discrete task (SD = 1.4 %) and 4.1 % of the rhythmic STOP trials (SD = 1.9 %).

### Measures

#### Reaction times (RT)

The behavioral results of these experiments have been published separately ^43^. Here and in the results section (below) we shortly present the behavioral measures that are essential to appreciate the main (EEG) results.

In the discrete experiment, RT_GO_ was calculated in the GO_D_ trials as the time between the primary stimulus onset and the response onset; the latter was defined as the moment the reach had exceeded 5 % of the Euclidean distance between the initial and furthest (i.e., end) position of the discrete-movement response. As an inhibitory RT, each participant’s RT_STOP-D_ was estimated using the integrative method for stop-signal tasks ^64,69^. In the rhythmic experiment, the movement-related StopTime was calculated as the time elapsed between the STOP signal onset and the end of the movement (i.e., null velocity). Each participant’s RT_STOP-R_, that is, the time between the STOP signal onset and the onset of movement alteration, was computed by identifying, within the StopTime, the first time point that the movement statistically deviated from the set of uninterrupted movements in the phase space ^4^.

#### Lateralized readiness potentials

In each condition LRPs were computed (using customized scripts written on Matlab) to assess the build-up of cortical motor activity following the primary stimulus (GO or CONTINUE). To this end, the EEG time series locked to the primary stimulus onset were averaged following the subtraction of a −200 to 0 ms pre-stimulus period as a baseline. The LRP was then derived from the difference between electrodes C3 (the electrode over the contralateral motor cortex) and C4 (its ipsilateral counterpart). This was done for GO_D_, successful-STOP_D,_ and fail-STOP_D_ trials in the discrete task and CONTINUE and STOP_R_ trials in the rhythmic one. As LRP is classically characterized by a negative deflection underlying motor preparation, LRP peak amplitude was defined in each condition by looking for the minimum peak value following stimulus onset (LRPs were 15 Hz low-pass filtered for the peak detection). A similar subtraction, that is, contralateral activity minus ipsilateral activity and vice versa, was performed for each pair of scalp electrodes (e.g., F3 minus F4, CP3 minus CP4 …) in order to display the lateralized part of the EEG activity as a topography (Fig. 1).

**Fig.1:**
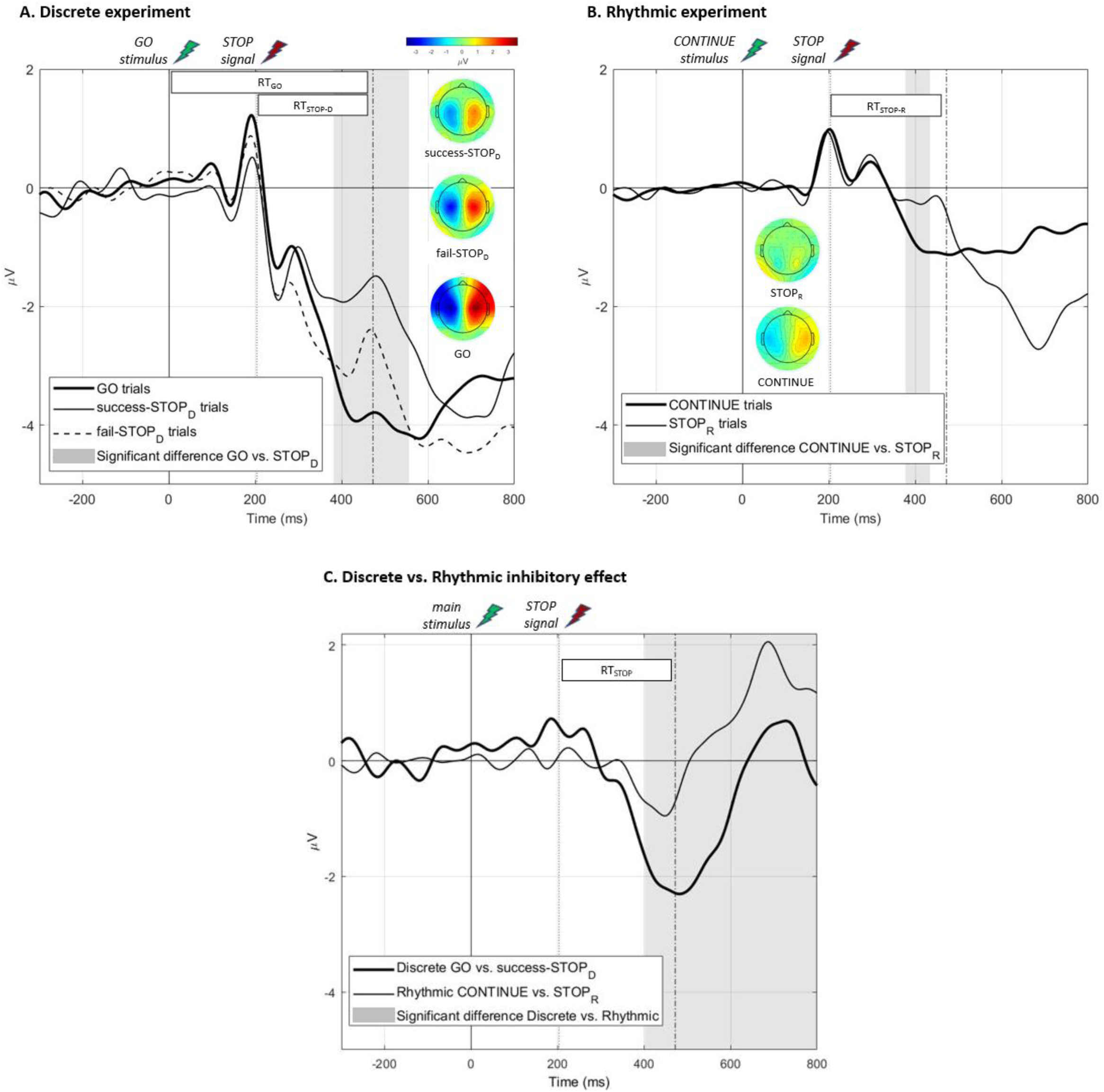
LRP analysis. **Panel A**: LRP (grand–average) computed in the discrete GO_D_, success-STOP_D_, and fail-STOP_D_ conditions. GO_D_ LRP differed significantly from success-STOP_D_ and fail-STOP_D_ conditions. In grey, the region of significant difference (according to the nonparametric permutation analysis) between GO_D_ and success-STOP_D_ conditions (p < .05, corrected). **Panel B**: LRP (grand–average) computed in the rhythmic CONTINUE and STOP_R_ conditions. In grey, the region of significant difference between the two conditions (p < .05, corrected). Topographies are presented in panels A and B as the lateralized topographies computed at each condition LRP peak latency (see Method section). **Panel C**: LRP inhibitory effect computed in the discrete (GO_D_ minus success-STOP_D_ LRP) and the rhythmic (CONTINUE minus STOP_R_ LRP) experiments. In grey, the region of significant difference between these two differential LRPs (p < .05, corrected). The represented SSD, RT_GO_ and RT_STOP_ latencies are based on the average of the obtained latencies over all the participants. LRPs were 15 Hz low-pass filtered for graphical purpose.

#### Mu and Beta time-frequency analysis

First, a time-frequency decomposition was performed according to the procedure described below, using the preprocessed EEG data from the C3 channel ^44,70,71^. The resulting time-frequency maps are shown for each experimental condition in Appendix B to provide a classical view of our data.

Second, a time-frequency analysis was performed with a focus on the Mu and Beta frequency bands. Thereto, the preprocessed EEG data were band-pass filtered in the 8 to 30 Hz frequency range. We then computed an ICA to this filtered data. This procedure of applying an ICA decomposition to a specific frequency-band is able to outperform the traditional wide-band ICA both in terms of signal-to-noise ratio of the separated sources and in terms of the number of the identified independent components ^72^. On the basis of the ICs resulting from the ICA algorithm, equivalent current dipoles were fitted using a four-shell spherical head model and standard electrode positions (DIPFIT toolbox ^73,74^). Then, to cluster ICs across participants, feature vectors were created combining differences in spectra (8−30 Hz), dipole location, and scalp topography. Clusters were next identified using a k-means clustering algorithm (*k* = 12) in EEGLAB. Among the resulting clusters, a single sensorimotor cluster was visually identified in each experiment (i.e., discrete and rhythmic) based on a centroparietal lateralized topography and a time-frequency map showing a clear ERD/ERS pattern.

In order to analyze the ERD/ERS activity of the MU and Beta bands, each IC of the two obtained clusters (i.e., discrete and rhythmic) was subjected to a time-frequency decomposition (using customized scripts written on Matlab) as follows: The EEG signals locked to the primary stimulus were convolved with complex 3-to-8 cycle-long Morlet’
ss wavelets. Their central frequencies were changed from 8 to 30 Hz in 0.5 Hz steps (and from 0.5 to 50 Hz for the C3 channel analysis in Appendix B). From the wavelet transformed signal, *w*_*k*_(*t, f*), of trial *k* at time *t* (3.5 ms time resolution) and with frequency *f*, the instantaneous power spectrum *p*_*k*_(*t, f*) = *R*(*w*_*k*_(*t, f*))^2^ + *I*(*w*_*k*_(*t, f*))^2^ was extracted (*R* and *I* symbolize the real and imaginary parts of a complex number, respectively). The mean power spectrum (i.e., averaged across trials) was then computed for each participant in the GO_D_, CONTINUE, STOP_D_, STOP_R_ and GO_R_ conditions as follow:

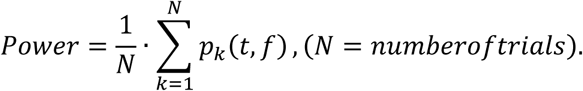

The power spectrum was then normalized with respect to a -400 to -100 ms pre-stimulus baseline and transformed to decibel scale (10 · log10 of the signal). In the rhythmic experiment, the baseline was extracted from the averaged GO_R_ trials (as in CONTINUE and STOP_R_ conditions, the pre-stimulus period includes movement). This mean power (time × frequency × power) was next averaged along the frequency dimension in an 8 Hz - 13 Hz window to compute the Mu power and a 15 Hz - 30 Hz window for the Beta power time series (time × power).

To detect significant ERD and ERS, the resulting Mu and Beta power time series of each condition was compared against the mean value of the power in the baseline time range (−400 to -100 ms). These comparisons were performed based on a non-parametric permutation procedure (see below). Thus, each time-period for which the power values were significantly below the baseline level was indexed as an ERD. Each time-period subsequent to an ERD and for which power did not significantly differ from the baseline level was indexed as an ERS. Each time-period including power values that were significantly above the baseline level was indexed as a power-rebound. To compare Mu and Beta dynamics between conditions, power time series were pairwise compared using the same non-parametric permutation procedure (see below).

#### Brain sources reconstruction

To estimate the brain structures pertaining to the clustered ICs, a brain-source reconstruction procedure was applied. For each clustered IC, the inverse ICA weight projections onto the original EEG channels were exported to the sLORETA (standardized low-resolution brain electromagnetic tomography) data processing module ^75^. sLORETA provides a unique solution to the inverse problem ^75–77^. For sLORETA, the intracerebral volume is partitioned into 6239 voxels with a 5 mm spatial resolution. Then, the standardized current density at each voxel is calculated in a realistic head model ^78^ based on the MNI152 template.

### Statistical analysis

To compare LRP time series between conditions at the group level, the LRPs were subjected to a nonparametric permutation procedure ^79^. Specifically, the 15 participants’ LRPs were pooled over the two compared conditions (15 per condition). Two sets of 15 LRPs were then drawn randomly (unpaired) from this pool, and the differential grand-average LRP was computed between the two sets. This procedure was repeated 10 000 times, thus producing a LRP distribution based on shuffled data under the null hypothesis. For each time point, a *p* value was computed as the proportion of these pseudo-differential LRPs that exceeded the observed participants’ average differential LRP. This *p* value indicates whether the observed power distribution for the two conditions diverged more than expected for random data (*p* = .05 threshold). To correct for multiple comparisons, we analyzed the resulting distributions of *p* values to compute *p* thresholds corresponding to the 2.5th percentile of the smallest, and the 97.5th percentile of the largest *p* values distribution ^80^. The same procedure was applied to the averaged Mu and Beta power time series to, first, assess ERD and ERS significance by comparing power time series against baseline values and, second, to asses power difference significance between conditions. In the case of the Mu and Beta power time series, the between-experiment comparison included an unequal number of ICs (20 discrete vs. 19 rhythmic ICs, respectively, see Results section). This variation was accounted for in the random-permutation stage of the statistical procedure by randomly selecting a pool of 19 ICs from each experimentation at each iteration.

Additionally, the study included measures of self-reported impulsivity, which were correlated with the EEG measures. This exploratory analysis was delegated to Appendix C for reasons of focus.

## Results

### Behavior

In the discrete experiment, the RT_GO_ (*M* = 472 ms, *SD* = 64 ms) and response probability (*M* = .54, *SD* = .08) permitted the estimation of individual’s RT_STOP-D_ (*M* = 269 ms, *SD* = 45 ms). The average STOP-signal delay (SSD) for participants was 203 ms (SD = 79 ms). In the rhythmic experiment, the spontaneous oscillation frequency was 1.65 Hz on average (SD = 0.54 Hz) and the analysis of the obtained StopTimes (*M* = 399, *SD* = 34 ms) enabled the computation of individual’s RT_STOP-R_ (*M* = 268, *SD* = 24 ms). Importantly, the RT_STOP-D_ and the RT_STOP-R_ values did not differ (*t* = .03, *p* > .05) and were unrelated across participants (*r* = .02, *p* > .05), suggesting independent but comparable timing of inhibition processing between the two experiments.

### Lateralized readiness potentials

In every condition, the LRP computation resulted in a typical negative deflection as portrayed in Fig. 1. In the discrete experiment, the permutation analysis identified a significant difference in the 381 - 556 ms time window (*p* < .05, corrected) between GO_D_ and successful-STOP_D_ conditions (Fig. 1.A) and in the 419 - 493 ms window between GO_D_ and fail-STOP_D_ conditions. In the rhythmic experiment, the same procedure identified a significant difference in the 377 - 434 ms time window (*p* < .05, corrected) between CONTINUE and STOP_R_ conditions (Fig. 1.B). To compare the “inhibitory effect” between the LRPs from the two experiments, differential LRPs were computed based on the GO_D_ minus successful-STOP_D_ difference for the discrete one and the CONTINUE minus STOP_R_ difference for the rhythmic one. The two differential LRPs were next compared through the same nonparametric permutation procedure, which revealed that the LRP reduction was significantly larger in the discrete experiment than in the rhythmic one in the 402 - 1,243 ms time window (*p* < .05, corrected; Fig. 1.C). Still, the peak amplitude of the differential LRP was significantly correlated between discrete and rhythmic experiments (*Pearson r* = .96, *p* < .001). Additionally, the exploratory analysis of individual’s motor impulsivity indicated a significantly lower LRP peak amplitude for the more impulsive participants in the GO_D_ and fail-STOP_D_ conditions (details in Appendix C.).

### Mu and Beta oscillations

The power maps resulting from the time-frequency decomposition applied to the preprocessed EEG data of the C3 channel (0.5 to 50 Hz) are shown for the different conditions in Appendix B.

In both experiments, only one sensorimotor cluster could be identified. Thus, a single sensorimotor cluster of 20 ICs (contribution of 15 participants) was retained for the discrete experiment. Another single cluster of 19 ICs was retained (15 participants) for the rhythmic experiment (Fig. 2.A). The power maps resulting from the time-frequency decomposition applied to the clustered components (8 to 30 Hz) are shown in Fig. 2.B. The detailed time course of Mu (8 - 13 Hz) and Beta (15 - 30 Hz) bands power and significant ERD/ERS are highlighted in Fig. 3. Overall, Mu and Beta power show the expected dynamics, that is, an ERD during the movement execution. This ERD appeared transient in the context of a discrete movement execution and sustained when the movement was rhythmic. The Mu/Beta ERD were followed by an ERS (Fig. 3). Notably, the ERS significantly exceeded the baseline level in the STOP_R_ condition only, evidencing of a post-movement Mu and Beta rebound in this condition.

**Fig.2:**
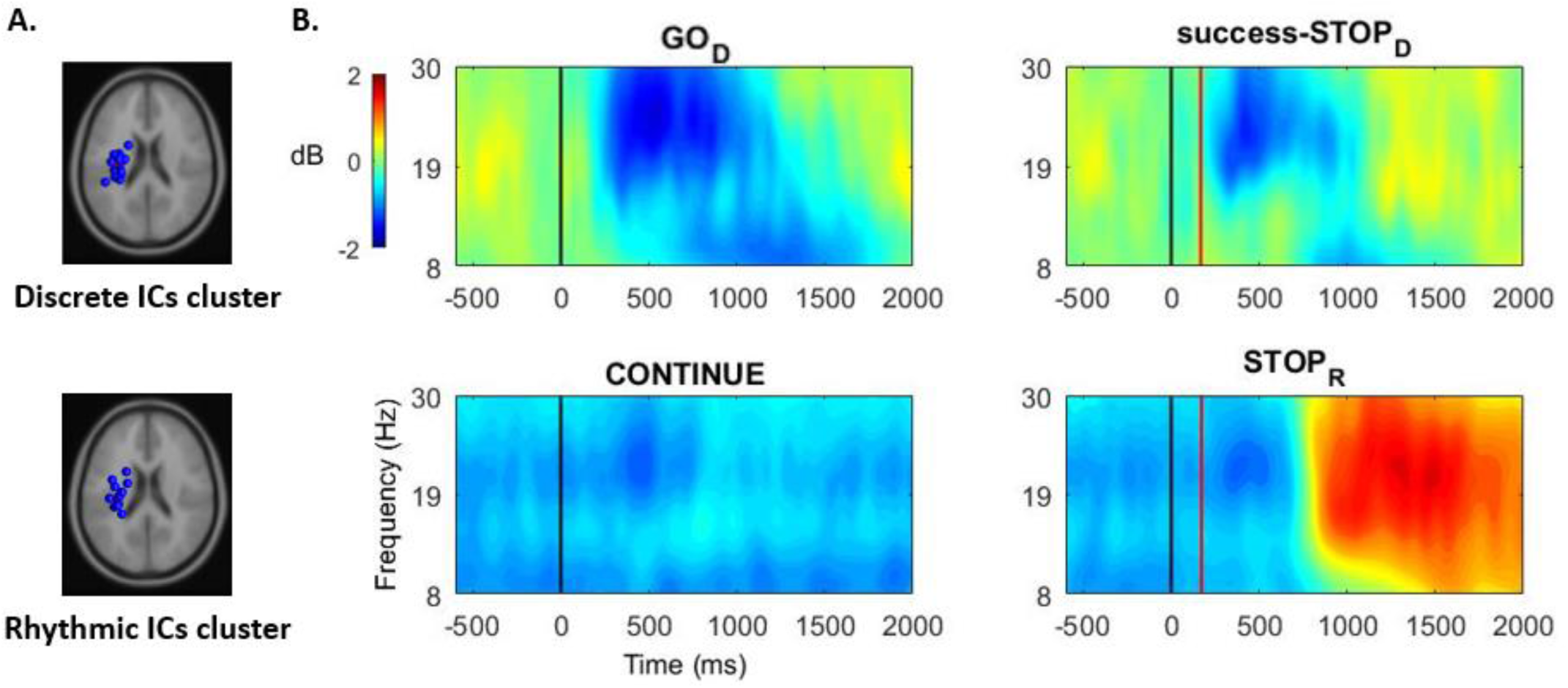
Component dimension time-frequency power analysis. **Panel A:** Equivalent current dipoles of the clustered sensorimotor components in the discrete (15 participants, 20 ICs) and the rhythmic (15 participants, 19 ICs) experiments. **Panel B:** Time-frequency power maps (ICs grand–average) computed in the discrete (GO_D_ and success-STOP_D_) and rhythmic (CONTINUE and STOP_R_) conditions. Black line: Primary (GO or CONTINUE) stimulus onset. Red line: STOP signal onset (the represented onset is based on the average of the obtained SSD, over all the participants). The blue scale represents desynchronization and the red scale (re)synchronization of the brain activity.

**Fig.3:**
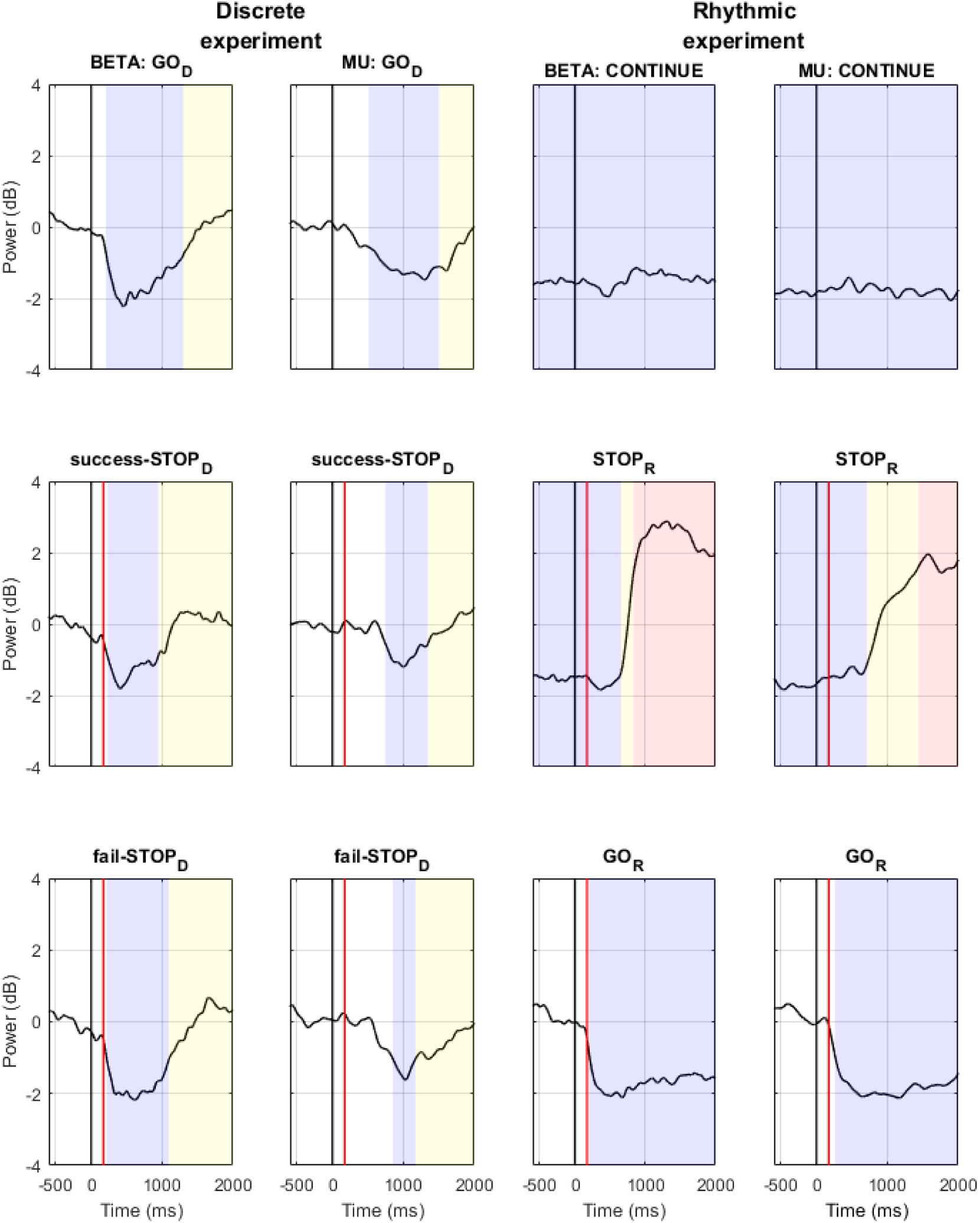
Beta and Mu power time series. Power time series (ICs grand–average) averaged in the Beta (15 to 30 Hz) and the Mu (8 to 13 Hz) frequency ranges from the time-frequency power maps computed in the discrete (GO_D_, success-STOP_D_ and fail-STOP_D_) and rhythmic (CONTINUE, STOP_R_ and GO_R_) conditions. Black line: Primary (GO or CONTINUE) stimulus onset. Red line: STOP signal onset (the represented onset is based on the average of the obtained SSD, over all the participants). Resulting from the non-parametric permutation comparison against baseline value, blue, yellow and red colors indicate time-ranges of significant ERD, ERS and power-rebound, respectively (see Method).

The Mu and Beta time-series were then compared between the experimental conditions in a pairwise fashion (non-parametric permutation procedure, see Method). The detailed result of these comparisons is provided in Table 1. Importantly, Mu power did not vary significantly between the three conditions of the discrete experiment: there was no significant difference the between movement-executed conditions (GO_D_ and fail-STOP_D_) and the no-actual-movement condition (success-STOP_D_). In the rhythmic experiment, the significantly higher Mu power in the STOP_R_ condition characterized a post-movement Mu ERS that was not present in the GO_R_ and CONTINUE conditions. When comparing the two experiments, the Mu power increase was stronger after the forced rhythmic-movement stop in the STOP_R_ condition as compared to all the other conditions, including the GO_D_ and success-STOP_D_ conditions, which are associated with a discrete-movement normal completion and cancellation, respectively.

**Table 1:** Pairwise condition comparison of Mu and Beta power time series. Mu and Beta power time series from the clustered ICs were compared between experimental conditions in a pairwise fashion using a non-parametric permutation procedure (see Method section). The resulting time-ranges of significant difference between conditions are reported. Z values indicate the threshold values corresponding to p < .05 (corrected for multiple comparisons, see Method) retained to assess significance. N.S. Non-significant.

| BETA power | GO<sub>D</sub> | success-STOP<sub>D</sub> | fail-STOP<sub>D</sub> | CONTINUE | STOP<sub>R</sub> | GO<sub>R</sub> |
| --- | --- | --- | --- | --- | --- | --- |
| MU power |  |  |  |  |  |  |
| GO<sub>D</sub> | – | Higher success-STOP<sub>D</sub> power from 1,161 to 1,287 ms<br>$z > 1.1473$ | N.S.<br>$z < 1.1911$ | Higher GO<sub>D</sub> power from -1,500 to 154 ms and from 1,468 to 2,000 ms<br>$z > 1.2438$ | Higher GO<sub>D</sub> power from -1,500 to 172 ms and higher STOP<sub>R</sub> power from 748 to 1,860 ms<br>$z > 1.7282$ | Higher GO<sub>D</sub> power from 1,374 to 2,000 ms<br>$z > 1.2392$ |
| success-STOP<sub>D</sub> | N.S.<br>$z < 1.5210$ | – | Higher success-STOP<sub>D</sub> power from 559 to 1,328 ms<br>$z > 1.1018$ | Higher success-STOP<sub>D</sub> power from -1,500 to 10 ms and from 1,133 to 2,000 ms<br>$z > 1.1908$ | Higher success-STOP<sub>D</sub> power from -1,500 to 154 ms and higher STOP<sub>R</sub> power from 780 to 2,000 ms<br>$z > 1.5789$ | Higher success-STOP<sub>D</sub> power from 1,091 to 2,000 ms<br>$z > 1.1954$ |
| fail-STOP<sub>D</sub> | N.S.<br>$z < 1.4692$ | N.S.<br>$z < 1.4689$ | – | Higher fail-STOP<sub>D</sub> power from -1,500 to 179 ms and from 1,447 to 2,000 ms<br>$z > 1.2893$ | Higher fail-STOP<sub>D</sub> power from -1,500 to 167 ms and higher STOP<sub>R</sub> power from 741 to 1,654 ms<br>$z > 1.7432$ | Higher fail-STOP<sub>D</sub> power from 1,325 to 2,000 ms<br>$z > 1.2455$ |
| CONTINUE | Higher GO<sub>D</sub> power from -1,500 to 397 ms and from 1,871 to 2,000 ms<br>$z > 1.6163$ | Higher success-STOP<sub>D</sub> power from -1,500 to 664 ms and from 1,458 to 2,000 ms<br>$z > 1.6883$ | Higher fail-STOP<sub>D</sub> power from -1,500 to 399 ms and from 1,804 to 2,000 ms<br>$z > 1.5852$ | – | Higher STOP<sub>R</sub> power from 773 to 2,000 ms<br>$z > 1.7528$ | Higher GO<sub>R</sub> power from -1,500 to 164 ms<br>$z > 1.1942$ |
| STOP<sub>R</sub> | Higher GO<sub>D</sub> power from -1,500 to 331 ms and higher STOP<sub>R</sub> power from 958 to 2,000 ms<br>$z > 1.7832$ | Higher success-STOP<sub>D</sub> power from -1,500 to 189 ms and higher STOP<sub>R</sub> power from 979 to 2,000 ms<br>$z > 1.7575$ | Higher fail-STOP<sub>D</sub> power from -1,500 to 175 ms and higher STOP<sub>R</sub> power from 928 to 2,000 ms<br>$z > 1.7198$ | Higher STOP<sub>R</sub> power from 888 to 2,000 ms<br>$z > 1.9067$ | – | Higher GO<sub>R</sub> power from -1,500 to -395 ms and higher STOP<sub>R</sub> power from 755 to 2,000 ms<br>$z > 1.8822$ |
| GO<sub>R</sub> | N.S.<br>$z < 1.5813$ | Higher success-STOP<sub>D</sub> power from 493 to 2,000 ms<br>$z > 1.6655$ | Higher fail-STOP<sub>D</sub> power from 1,152 to 2,000 ms<br>$z > 1.5310$ | Higher GO<sub>R</sub> power from -1,500 to 178 ms<br>$z > 1.6104$ | Higher GO<sub>R</sub> power from -1,500 to -216 ms and higher STOP<sub>R</sub> power from 865 to 2,000 ms<br>$z > 1.9595$ | – |

Regarding the Beta power, the discrete conditions GO_D_ and fail-STOP_D_ in which the movement was executed did not significantly differ. In contrast, the success-STOP_D_ condition exposed a higher Beta power than GO_D_, from 1,161 to 1,287 ms, and than fail-STOP_D_, from 559 to 1,328 ms. In the rhythmic experiment, the significantly higher Beta power in the STOP_R_ condition related to a post-movement Beta ERS that was not present in the GO_R_ and CONTINUE conditions (Fig. 3.). When comparing the two experiments, the pattern of differences was similar to the Mu power, with the post-movement Beta power increase being stronger in the STOP_R_ than the GO_D_ or the success-STOP_D_. Additionally, the exploratory analysis of individual’s motor impulsivity indicated significantly a higher PMBR amplitude for the more impulsive participants in the STOP_R_ conditions (details in Appendix C.).

### Brain sources reconstruction

Based on the voxel-based sLORETA images, we searched for brain activation using voxel-wise randomization *t*-tests with 5000 permutations, based on nonparametric statistical mapping. This procedure was performed separately for the ICs of the discrete and rhythmic clusters. Significant voxels (*p* < .01, corrected for multiple comparisons) were located in the MNI-brain (Fig. 4) regarding the engaged Brodmann areas (BA) and the voxels coordinates. In the discrete experiment, the clustered ICs activity was related to the activation of sensory regions such as the primary somatosensory (BA 1, BA 2, BA 3) and the somatosensory association (BA 5) cortices, as well as M1 (BA 4). In the rhythmic experiment, activation was found in the primary somatosensory cortex (BA 3), as well as pre-motor areas (BA 6), and M1 (BA 4) (detailed MNI coordinates of the activation are provided in Table 2).

**Fig.4:**
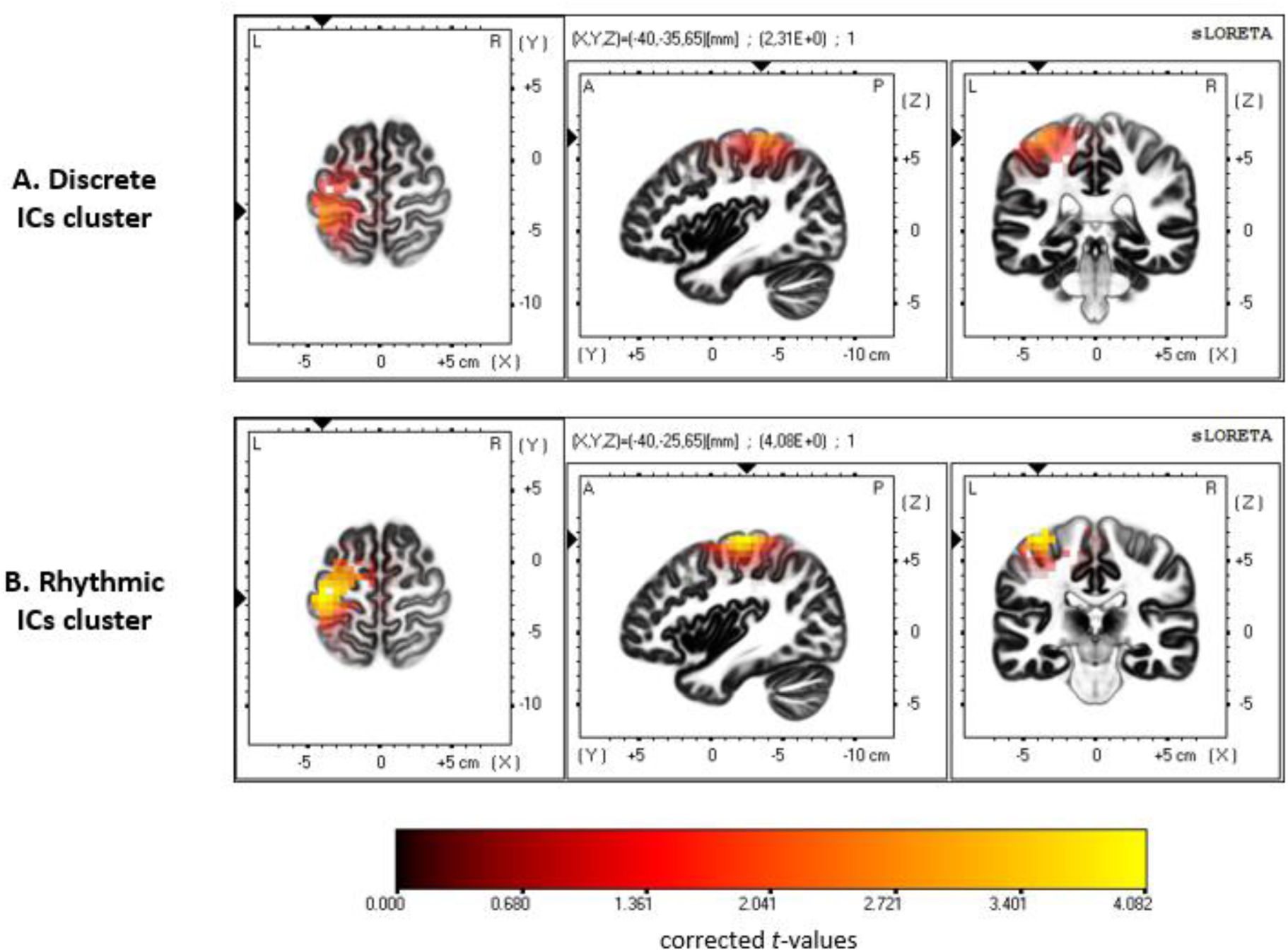
Brain sources reconstruction. The sLORETA images showing significant estimated activation pertaining to the discrete (**panel A**) and rhythmic (**panel B**) clustered ICs, for three orthogonal brain slices (horizontal, sagittal, coronal). Only the voxels that passed the p value threshold (p < .01, corrected) are shown in color. The color represents t value. In the discrete experiment, activation was found in sensory (BA 1, BA 2, BA 3, BA 5) and motor areas (BA 4). In the rhythmic experiment, fewer sensory (BA 3) but (one) more motor regions (BA 4, BA 6) were involved. Detailed MNI localization of the significant activation is provided in Tab. 2.

**Tab. 2:**
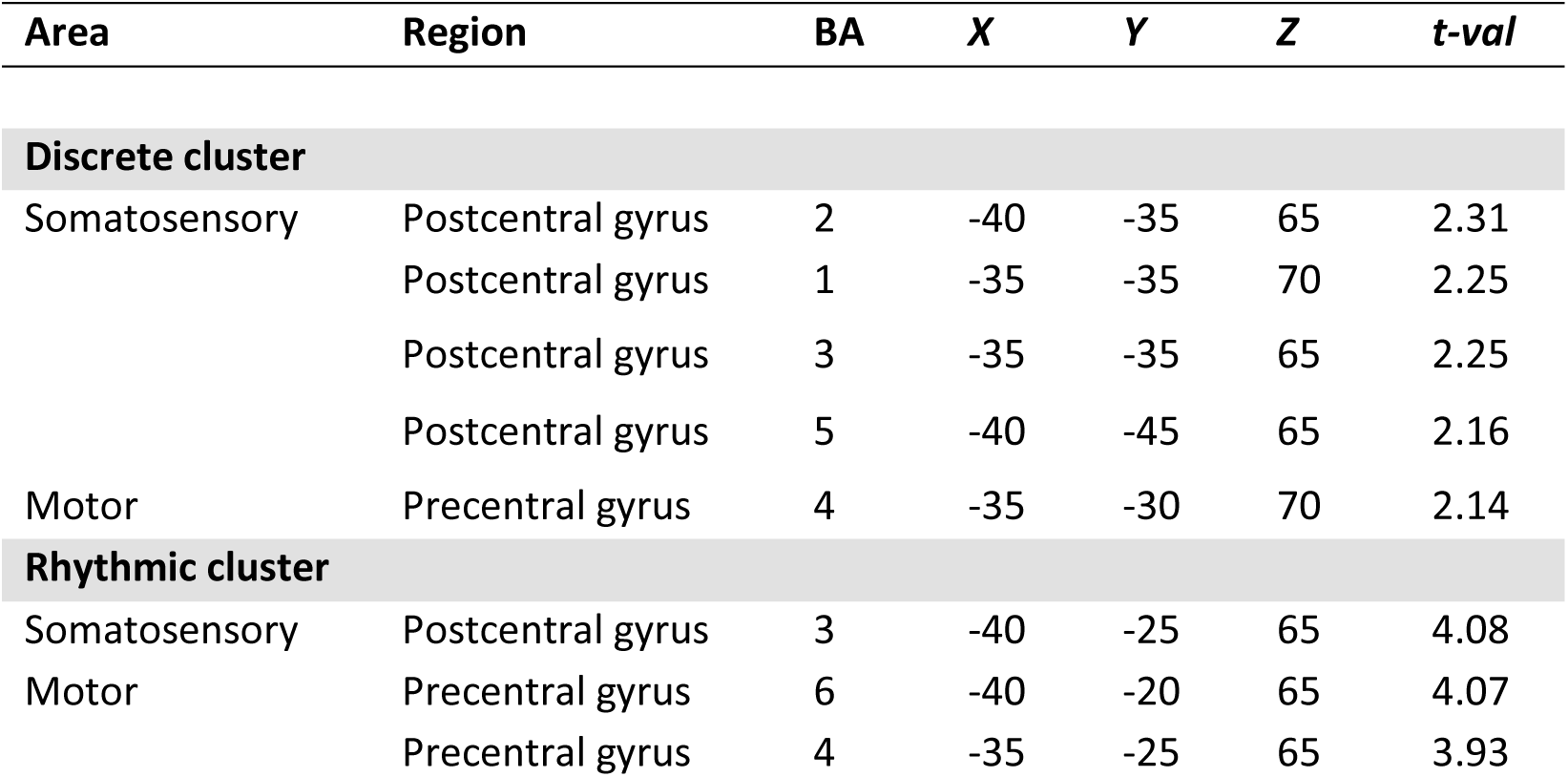
Summary of significant activation from the brain sLORETA reconstruction. Significant (p < .01, corrected) regions are indicated with the name of Brodmann area (BA), MNI coordinates (X, Y, Z) and t value (t-val) of the higher statistical tresholded voxel.

## Discussion

The present study examined the neural sensorimotor activity related to performing and suppressing movements pertaining to the discrete and rhythmic classes. EEG data were analyzed in both contexts to provide new insight into the function of LRP and sensorimotor ERD/ERS patterns in the Mu and Beta frequency bands. Notably, the estimated generators of the cortical ERD/ERS pattern identified over peri-Rolandic areas closely overlap those reported in previous work ^14,50,52,62^. Indeed, we identified both somatosensory and motor cortical areas as generators of the observed ERD/ERS pattern, supporting the idea that both movement-related and sensory-related neural activity may be engaged. The inhibition mechanism triggered by the STOP signal affected the LRP in both the discrete and rhythmic experiments, and occurred before the end of the RT_GO_, RT_STOP-D_, or RT_STOP-R_ latencies. Additionally, the measured RT_GO_ for movement generation and RT_STOP_ for movement suppression fell in the time range classically observed in stop-signal experiments across various movement responses ^81–84^. The similarity between the discrete and the rhythmic RT_STOP_ values indicates that the processes engaged in aborting the two movement classes are of comparable duration.

Our first expectation dealt with the LRP dynamics. We hypothesized a large LRP following a GO stimulus to contrast with the absence of an LRP (i.e., zero amplitude) following a CONTINUE stimulus, and that this LRP amplitude would be reduced by the STOP signal occurrence only in the discrete experiment. For the discrete movements, an LRP was triggered by the primary GO stimulus, and was subsequently impacted by the STOP signal in both successful-STOP_D_ and failed-STOP_D_ trials. These findings are consistent with the notion that an inhibition signal that arrives at M1 attenuates cortical motor outflow, as reflected by the reduction of the LRP amplitude ^40^; in the case of fail-STOP_D_ trials, this reduction is insufficient to restraint the response threshold to be reached ^33^. For the rhythmic movements, the CONTINUE stimuli occurring during the ongoing movement also led to an LRP response, albeit weaker than in the GO_D_ instruction. Rebutting our hypothesis, this LRP response indicates that the presentation of the CONTINUE stimulus during the ongoing movement triggers a non-negligible cortical motor activity. Thus, LRP might not index pre-movement processing only, but also any cortical motor activity occurring before and during movement. Alternatively, if the rhythmic movement is implemented as a concatenation of discrete units, the LRP might reflect the cortical motor activity engaged in the initiation of each unit.

Indeed, previous studies have shown that the sensorimotor activity recorded in rhythmic movements suggested a discrete-units-concatenation when the movement frequency was ranging from 0.33 to 1 Hz, whereas this activity was ‘truly’ continuous for above 1 Hz movement frequencies ^31,32^. Nevertheless, as the rhythmic movements in the present study were, on average, performed at 1.65 Hz, the LRP observed in the CONTINUE trials are unlikely to reflect motor cortical activity related to the concatenation of discrete movements.

The LRP amplitude following the CONTINUE stimulus was reduced in the STOP_R_ condition. Notably, the amplitude of this “inhibitory effect”, albeit weaker, was strongly correlated to the GO minus STOP_D_ LRP difference measured in the discrete experiment. Thus, the LRP reduction might index action inhibition in the context of both prepared-discrete and ongoing-rhythmic movement suppression. This interpretation is consistent with the notion that LRP is a marker of the cortical motor activity as a common final pathway in the central control of movement and thus be the “site” where (frontal) executive “agents” exert inhibitory control ^85^. Note that the commonality of the motor site of inhibition in discrete and rhythmic action inhibition does not provide information about the inhibiting “agents” engaged in the two situations, as the two levels of inhibition processing can be independent ^40^. Notably, the EEG markers of the executive agents engaged in action inhibition tended to dissociate the processing of discrete action cancelling and rhythmic action stopping ^43^.

Our second expectation that the Mu ERD/ERS observed pattern should show a transient vs. sustained activity for the discrete and rhythmic experiments, respectively, was confirmed. This validates the discrete vs. rhythmic nature of the performed movements and aligns with the understanding of the Mu rhythm as a correlate of the interaction between sensory and motor information processing: The sustained ERD during ongoing movement may correspond to a closed-loop control for the online control of the ongoing movement. In contrast, the transient Mu ERD/ERS pattern did not differ between the performed discrete actions in the GO_D_ and fail-STOP_D_ conditions and the cancelled ones in the success-STOP_D_ condition. This finding is in line with the Mu rhythm being independent of the movement outcome, which may be the case if the Mu rhythm encodes the processing of sensorimotor integration in an open-loop control of discrete actions. Notably, the reactive inhibition of discrete actions has been coherently conceptualized as a dual-step process encompassing attention reorientation (by the STOP signal) and prepared-movement cancellation ^86–88^. The reorientation of attention is not specific to action inhibition but generalizes to multiple situations implicating goal redirection, including the reaction to a GO stimulus ^88^. Following the hypothesis that the Mu rhythm is an alpha-like oscillation that links perception and action ^49,50^, a Mu ERD is expected to occur when cortical motor activity is modulated following attentional reorientation, which includes both discrete GO_D_ and STOP_D_ trials. Hence, the absence of actual movement in successful-STOP_D_ trials should not modulate the Mu rhythm dynamics relative to Mu rhythm in GO_D_ trials. Our results are in accordance with this expectation. A compatible finding is that the Mu ERD/ERS varies with attention ^89^.

Confirming our third expectation, the Beta ERD appeared sustained for the ongoing rhythmic movement whereas it was transient for the discrete movement, thus following the motor activation dynamics. Next, a Beta ERS occurred following the action. This is consistent with the purported role of the Beta ERS in evaluating the action sensory output, in that it was lower for discrete-movement failed cancellation compared to successful cancellation. Previous findings already reported this “error-related Beta rebound reduction”, which may relate to salient error/mismatch detection mechanisms ^90,91^. Still, some results diverged from our expectations. On the one hand, the PMBR was higher following a forced rhythmic movement stop (STOP_R_) than the Beta ERS following discrete movement completion (GO_D_). On the other hand, the discrete-action Beta ERS was higher after a successful action cancellation than following action completion in GO_D_ and fail-STOP_D_ conditions, which did not differ in this regard. These two findings support the notion that a higher Beta ERS is a correlate of active action suppression ^14^, here triggered by a STOP signal. Whereas Parkes et al. identified PMBR neural generators in post-Rolandic (sensory) areas, which they interpreted in favor of the notion that PMBR reflects sensory reafference evaluation ^59^, other studies suggested that the PMBR was also related to pre-Rolandic (motor) activation ^58,92,93^. Our results are in line with the latter findings, with both a significant PMBR and pre-motor activation being reported for the rhythmic but not discrete actions. This engagement of pre-motor cortices in the rhythmic movements is congruent with the previously reported pre-supplementary motor area activation in PMBR ^94,95^. Thus, our results do not exclude that Beta ERS is an index of action sensory outcome evaluation, but they also support the view that it is associated with an active inhibition process of cortical motor activity.

Nonetheless, this active inhibition hypothesis of the PMBR functional role is silent on why the ongoing action-forced stop gave rise to a large PMBR over contralateral sensorimotor cortical areas, whereas a much weaker Beta ERS followed discrete action cancellation. A tentative explanation is that the inhibitory process engaged in movement cancellation acts at the movement preparation level, as indicated by the LRP decrease and the ERD abortion in the STOP_D_ condition ^41^. Thus, inhibition might lie in maintaining the cortical idle state to cancel a discrete action, whereas it would force the return to this idle state to stop a rhythmic movement. This explanation is also consistent with the notion that a discrete action, if controlled in an open-loop fashion, is not associated with an online control based on sensory prediction evaluation, as the PMBR is a correlate of the latter. In contrast, if controlled in a closed-loop fashion, the ongoing-rhythmic action requires the evaluation of the sensory predictions associated with the movement production, as indicated by a significant PMBR. A distinction in the movement-suppression after-effect (i.e., PMBR) suggests that discrete-action cancelling and rhythmic-action stopping may engage distinct inhibition processes ^43^. As action inhibition operates on both discrete ^64,81^ and rhythmic ^4–6,42^ movements, considering the distinction between the two movement classes would undoubtedly contribute to a better understanding of this complex process at the neurobiological level.

Alternatively, the lower Beta ERS following discrete action completion and cancellation compared to the large PMBR following rhythmic action stop, may reflect a PMBR that has been reduced due to the task uncertainty. Indeed, previous work suggested that beta power reflects the estimated uncertainty in the parameters of the forward models involved in motor control ^96^. Thus, the primary stimuli (blue or green) in the discrete experiment required a two-choice reaction (i.e., trigger a discrete movement toward the left or right side), whereas the same stimuli required a unique response in the rhythmic experiment (i.e., continue the movement for both blue and green stimuli). This discrepancy may introduce a modulation of confidence in the predicted sensory outcome in the forward model of action control, resulting in a lower post-movement Beta modulation ^54,96^. In contrast, the rhythmicity of an ongoing movement may lead to a confident movement execution that increases the PMBR ^97^.

Overall, our pattern of results regarding the Beta power dynamics excludes an understanding of the PMBR neither as a correlate of the action sensory outcome evaluation nor as an index of active motor suppression. In fact, both interpretations are not incompatible, and a tentative explanation is that the PMBR reflects the action control in forward models, with its amplitude being modulated by the uncertainty and the engagement of an inhibition process. Thus, the PMBR could be reduced when the uncertainty of the predicted sensory output is high, whereas it would be strengthened in reaction to an inhibition signal. This imperative action suppression might result in suppressing the motor plan execution and its predicted sensory outcome. It could also lead to the interruption of the closed-loop processing of sensorimotor information itself, as indicated by the Mu rebound that followed the rhythmic action stop. Although this explanation remains highly hypothetical without studies manipulating sensory feedback and inhibition requirement, it globally fits well with a recently established framework in which Beta rebounds reflect, at various cortical sites, a “clearing-out” of the motor plan ^98^.

The present study focused on the movement performance and suppression in reaction to an external cue, so-called exogenous action control ^99^. Adapted behavior also includes performing and suppressing movement in a self-initiated fashion, that is, endogenous motor control. Generalizing the present functional interpretation of neural sensorimotor activities requires that future experiments study and contrast both situations. Especially, internal and external movement initiation require partially distinct sensorimotor activities ^100^. Movement suppression mechanisms are also known to vary as a function of whether proactive vs. reactive inhibition is required, both for the suppression of discrete ^101,102^ and rhythmic ^6^ movements. These investigations are much needed to provide a complete comprehension of sensorimotor cortical activity.

The understanding of sensorimotor activity has implications for multiple clinical syndromes associated with movement disorders ^103^. The abilities to initiate and stop action are especially affected by impulsivity ^3^, which is an essential dimension of several psychiatric disorders, such as attention deficit hyperactivity disorder (ADHD) and obsessive-compulsive disorder (OCD). Evaluating neural sensorimotor activity through movement-related cortical ERD/ERS, healthy participants have been distinguished from those with ADHD ^104^ and OCD ^105^. In the general population, sensorimotor activity is poorly investigated in relation to individuals’ impulsivity traits. A recent study suggested that sensorimotor ERD/ERS amplitude may relate to impulsivity ^106^. The association reported in Appendix C. between motor impulsivity and lower LRP amplitude in triggering a discrete action and higher PMBR when forced-stopping a rhythmic action suggests that cortical sensorimotor activity in the execution and suppression of action might depend on the individual’s impulsivity level. Still, further studies targeting the impulsivity dimension and including participants exhibiting a broad range of impulsivity levels are required to test this hypothesis.

Finally, the present study provides new insights in understanding the cerebral sensorimotor activity by exploring EEG records of LRP and Mu/Beta rhythms associated with the performance and suppression of movement in the context of discrete and rhythmic classes of actions. Showing the distinct sensorimotor dynamics that operate in the two action classes, our findings are highly compatible with recent proposals that Mu and Beta rhythms might encode reciprocal interactions between motor and sensory cortices to enable movement monitoring ^47,96^. Still, the PMBR may also reflect the engagement of a clearing-out function to abort the sensorimotor processing when action has to be inhibited ^14^. At any rate, our findings support the notion that Mu and Beta frequency bands play complementary roles in the sensorimotor control of action. Further studies using imaging procedures with a better spatial resolution are required to disentangle the Mu and Beta specific implication in the different cortical areas that engage in action performance and suppression.

## Data Availability

The data generated during and/or analyzed during the current study are available from the corresponding author upon reasonable request.

