## Appendices for "Cortical sensorimotor activity in the execution and suppression of discrete and rhythmic movements"

### Appendix A.

#### Experimental procedures

Participants performed two movement experiments: a discrete one and a rhythmic one. Both experiments were performed on the same WACOM Cintiq 15X tablet (1280×800-pixel resolution). As long as the stylus touched the tablet, the x and y coordinates of the performed motions were digitized at a sampling frequency of 143 Hz. The program controlling the tablet was custom-made.

In the discrete experiment, participants adopted, as an initial state, a static position (i.e., no movement), which consisted of keeping the stylus between two vertical yellow bands (1 mm wide) plotted at the center of the digitizing black screen (10 mm distant). In the rhythmic experiment, participants were instructed to continuously oscillate at a spontaneous frequency with the stylus between the two sides of the screen but with the oscillation extrema falling outside the two centered vertical lines. In this experiment, the initial state is then a rhythmic movement, thus corresponded to 'being in movement'.

To drive a primary (perform) task, stimuli were green or blue 50 ms flashes displayed on the whole screen. In the discrete experiment, participants were instructed to reach with the stylus to the right versus left half-side of the tablet screen when a green versus blue flash appeared, respectively (green and blue stickers were visible on the right and left tablet sides; **Supplementary Fig. 1**). Therefore, the primary discrete task consisted of a two-choice reaction time involving a discrete-action response ($GO_D$ condition). In the rhythmic experiment, the primary task was to pursue the rhythmic movement without interruption when the green and blue stimuli appeared (CONTINUE condition).

As a secondary (stop) task, in 25 % of the trials, the primary-task stimulus was followed by a red 50 ms flash, which indicated to the participants to abort their primary-task response. Thus, they had to either cancel the prepared-discrete movement ($STOP_D$ condition) or stop the ongoing-rhythmic movement ($STOP_R$ condition). In the rhythmic task, a $GO_R$ trial was added after each STOP trial to re-engage participants in the rhythmic movement. In the discrete experiment, the stop-signal delay between the primary-task stimulus and the stop signal (SSD), initially set to 200 ms, was dynamically adjusted in 50 ms increments to achieve a probability of responding p(respond|signal) of .50. When the participant crossed a vertical line, the STOP trial was considered as a stop failure and the SSD was shortened; when the participant kept the stylus between the two lines, the STOP trial was considered successful and the SSD was prolonged. In the rhythmic experiment, the SSD value was set to a fixed value equivalent to the mean of the SSD obtained by each participant in the discrete experiment. To this purpose, all participants completed the discrete experiment one week prior to the rhythmic one.

In both experiments, the participants completed one practice block and 30 experimental blocks, each consisting of 20 trials. The discrete experiment was set up according to the standard guideline for stop-signal experiments (Verbruggen et al., 2019). The rhythmic experiment was built symmetrically to the discrete one, in order to limit the difference between the two experiments to the modality of the movement involved in the primary task (i.e., discrete or rhythmic).

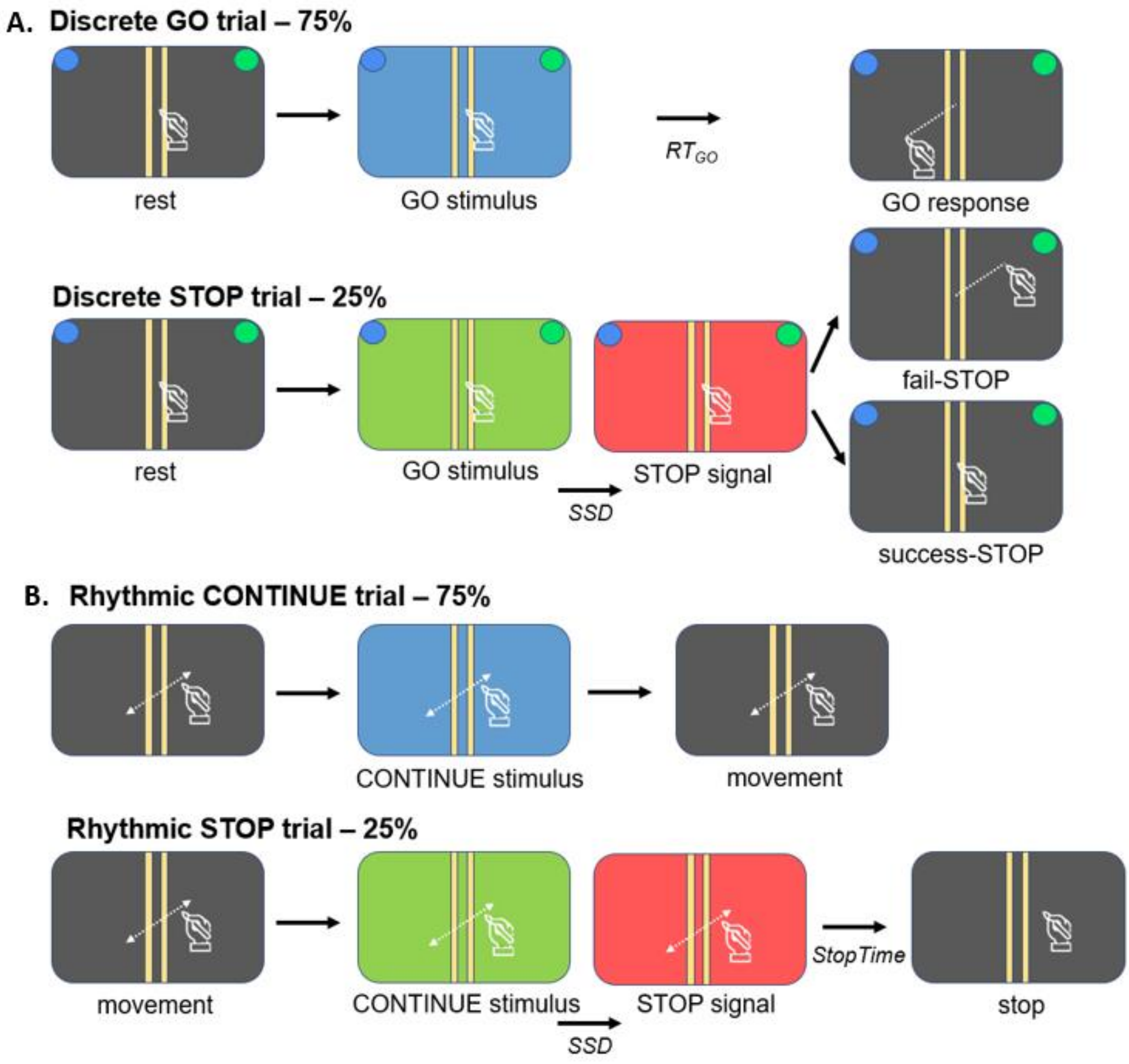


**Supplementary Fig. 1**: **Discrete and rhythmic experimental designs**

*Participants responded to the primary stimulus by initiating a discrete movement (in the discrete GO condition; panel A) or continuing a rhythmic movement (in the rhythmic CONTINUE condition; panel B). In 25 % of the trials, the primary stimulus was followed by a STOP signal after a SSD, which was variable in the discrete task but fixed in the rhythmic one (see text).*

**Appendix B.**

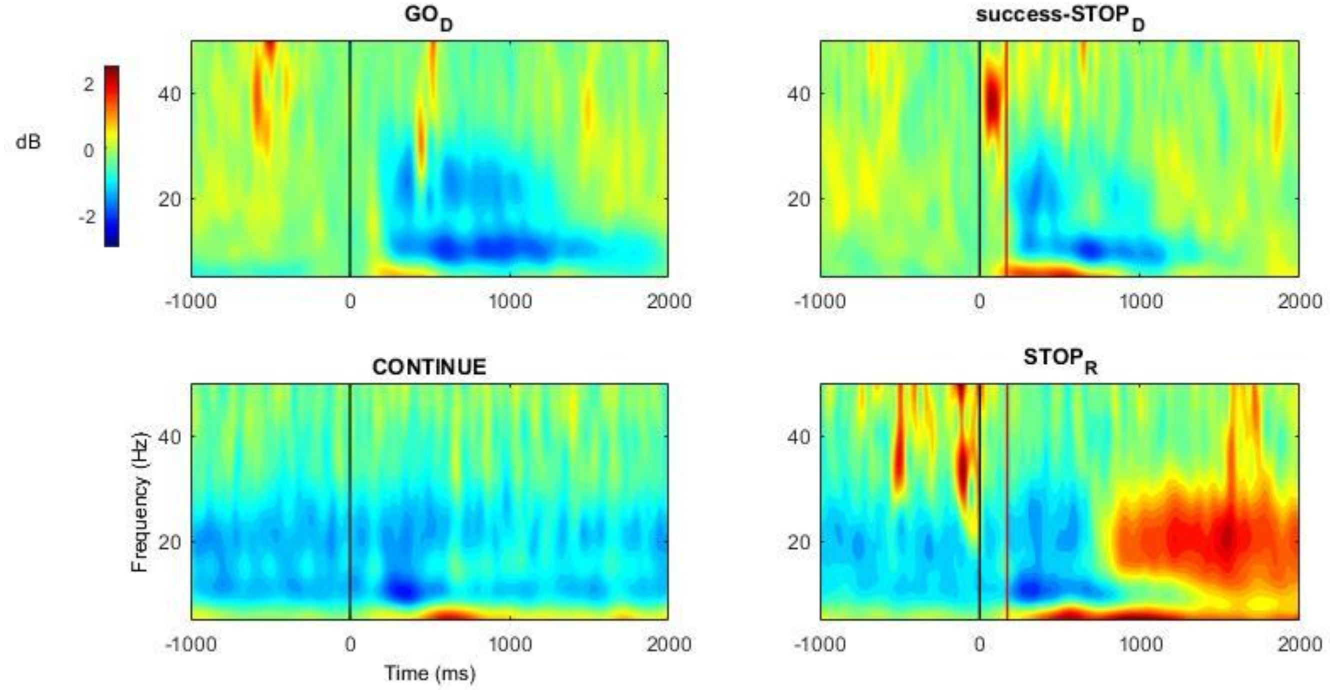


**Supplementary Fig. 2**: **C3 channel time-frequency power analysis**

*Time-frequency power maps (grand–average) computed from the C3 channel in the discrete ($GO_D$ and success-$STOP_D$) and rhythmic (CONTINUE and $STOP_R$) conditions. Black line: Primary (GO or CONTINUE) stimulus onset. Red line: STOP signal onset. The blue scale indicates a desynchronization and the red scale a (re)synchronization of the neural activity.*

**Appendix C.**

### Self-reported impulsivity

During the first session of the two experimental sessions completed by the participants of the study, the Barratt Impulsiveness Scale (BIS) questionnaire was adopted to assess the behavioral traits of impulsiveness for each participant (Patton et al., 1995). Using a French version of the questionnaire (Baylé et al., 2000), the items were rated on a four-point Likert scale from "never" to "almost always/always", in which higher scores indicate higher levels of impulsivity. The BIS questionnaire is subdivided into three sub-dimensions. Non-planning ($BIS_{PLANNING}$) is evaluated as a tendency to plan and think carelessly, cognitive impulsivity ($BIS_{COGNITION}$) refers to difficulties in focusing on a task, and motor impulsiveness ($BIS_{MOTOR}$) is a tendency to act on the spur of the moment.

Each participant's average impulsivity score was computed for the overall BIS questionnaire ($M$ = 62.9, $SD$ = 8.1) as well as for the single $BIS_{MOTOR}$ ($M$ = 22.1, $SD$ = 3.5), $BIS_{COGNITION}$ ($M$ = 17.7, $SD$ = 3.8), and $BIS_{PLANNING}$ ($M$ = 23.1, $SD$ = 3.5) components.

Pearson correlations were performed to evaluate the linear relation between the individual's impulsivity scores and EEG measures (LRP and ERD/ERS peak amplitudes). The obtained $p$ values were adjusted according to the number of computed correlations using a Holm-Bonferroni procedure.

Significant correlations were found between $BIS_{MOTOR}$ and LRP peak amplitude of the $GO_D$ ($r$ = .56, $p$ < .05, corrected) and fail-$STOP_D$ ($r$ = .55, $p$ < .05, corrected) conditions in the discrete experiment, indicating lower LRP amplitude for more impulsive participants. No correlation was found with LRP values in the rhythmic experiment. No correlation between participants' scores and the Mu peak ERD/ERS amplitude reached significance following the correction for multiple tests regarding the self-reported impulsivity. A fairly strong correlation was found between $BIS_{MOTOR}$ and the Beta ERS of the $STOP_R$ condition ($r$ = .72, $p$ < .05, corrected), showing an association between larger PMBR and higher impulsivity scores.

Baylé, F. J., Bourdel, M. C., Caci, H., Gorwood, P., Chignon, J.-M., Adés, J., & Lôo, H. (2000). Structure factorielle de la traduction française de l'échelle d'impulsivité de Barratt (BIS-10). *The Canadian Journal of Psychiatry*, *45*(2), 156-165. https://doi.org/10.1177/070674370004500206

Patton, J. H., Stanford, M. S., & Barratt, E. S. (1995). Factor structure of the Barratt impulsiveness scale. *Journal of Clinical Psychology*, *51*(6), 768-774. https://doi.org/10.1002/1097-4679(199511)51:6<768::aid-jclp2270510607>3.0.co;2-1

Verbruggen, F., Aron, A. R., Band, G. P., Beste, C., Bissett, P. G., Brockett, A. T., Brown, J. W., Chamberlain, S. R., Chambers, C. D., Colonius, H., Colzato, L. S., Corneil, B. D., Coxon, J. P., Dupuis, A., Eagle, D. M., Garavan, H., Greenhouse, I., Heathcote, A., Huster, R. J., … Boehler, C. N. (2019). A consensus guide to capturing the ability to inhibit actions and impulsive behaviors in the stop-signal task. *eLife*, *8*, e46323. https://doi.org/10.7554/eLife.46323
